# Mechanisms of action and synergies of a novel lipid IV_A_ biosynthesis inhibitor

**DOI:** 10.1101/2023.09.15.557861

**Authors:** Emma R Holden, Muhammad Yasir, A Keith Turner, Mark A Webber, Ian Charles, Ed Siegwart, Tony Raynham, Ajay Mistry, John George, Matthew Gilmour

## Abstract

The development of novel antimicrobials provides additional treatment options for infectious diseases, including antimicrobial resistant infections. There are many hurdles to antimicrobial development and identifying an antimicrobial’s mechanism of action is a crucial step in progressing candidate molecules through the drug discovery pipeline. We used the genome wide screening method TraDIS-*Xpress* to identify genes in two model Gram-negative bacteria that affected sensitivity to three analogues of a novel antimicrobial compound (OPT-2U1). TraDIS-*Xpress* identified that all three analogues targeted the lipid IV_A_ biosynthetic pathway in *E. coli* and *Salmonella* Typhimurium. Specifically, we determined that the antimicrobial target was likely to be LpxD, and validated this by finding a 5 log_2_-fold increase in the MIC of the OPT-2U1 analogues in *E. coli* when *lpxD* was overexpressed. Synergies were identified between OPT-2U1 analogues combined with rifampicin or colistin, to varying strengths, in both *E. coli* and *S*. Typhimurium. LPS composition was a likely reason for differences between *E. coli* and S.Typhimurium, as perturbation of LPS synthesis affected synergy between antibiotics and OPT-2U1 analogues. Finally, genes involved in ATP synthesis and membrane signalling functions were also found to affect the synergy between colistin and OPT-2U1 analogues. TraDIS-*Xpress* has proven a powerful tool to rapidly assay all genes (and notably, essential genes) within a bacterium for roles in dictating antimicrobial sensitivity. This study has confirmed the predicted target pathway of OPT-2U1 and identified synergies which could be investigated for development of novel antimicrobial formulations.

**Data Summary:** Nucleotide sequence data supporting the analysis in this study has been deposited in ArrayExpress under the accession number E-MTAB-13250. The authors confirm all supporting data, code and protocols have been provided within the article or through supplementary data files.

## Introduction

Infections caused by antimicrobial-resistant pathogens are increasing in incidence and severity around the world (Cassini et al., 2019). By 2050, approximately 10 million people per year are predicted to die due to complications associated with antimicrobial-resistance (O’Neill, 2016). Interventions are needed to reduce the development of antimicrobial-resistant pathogens and lower the burden of disease. These may include the development of new antimicrobials with novel mechanisms of action. Novel antimicrobials, used either on their own or synergistically with other antimicrobials, have the potential to reduce the development of antimicrobial resistance and the incidence of treatment failure. However, the development of novel antimicrobial agents has many economic and biological hurdles that must be overcome before clinical trials can commence.

Identifying the mechanism of action of novel antimicrobials is essential for their continuation through the drug development pipeline. Recently, the massively parallel transposon mutant screen TraDIS-*Xpress* was used to assay every *E. coli* gene to identify those involved in survival under antimicrobial stress, including to triclosan (Yasir et al., 2020), fosfomycin (Turner et al., 2020), meropenem (Thomson et al., 2022) and colistin (Yasir et al., 2022). This methodology identified known and unknown antibacterial susceptibility mechanisms in *E. coli* and predicted mechanisms of synergy between trimethoprim and sulfamethoxazole (Turner et al., 2021). Notably, the very large transposon mutant libraries make use of a transposon-encoded outwards-transcribing inducible promoter, allowing investigation into how gene expression, as well as gene disruption, affects survival under a growth condition of interest. The modulation of expression in neighbouring genes also allows investigation of essential genes, which is of utmost importance given many antimicrobials often target essential genes.

We used TraDIS-*Xpress* to determine the mechanism of action of a series of analogues of a novel antimicrobial compound OPT-2U1. This compound was discovered by screening compounds for the potentiation of rifampicin and structure-based design was applied to strong potentiators yielding direct acting molecules (analogues named 003, 009 and 010, structure not shown pending confidential intellectual property). We measured transposon mutant susceptibility to these analogues in both *Escherichia coli* and *Salmonella enterica* serovar Typhimurium and confirmed the compounds likely affect LPS biosynthesis through inhibiting the synthesis of lipid IV_A_. LPS may prove to be a useful target for antibiotic development as pathways are fairly conserved across Gram-negative bacteria (Sabnis and Edwards, 2023). We also identified and described the mechanisms behind a synergistic relationship between OPT-2U1 and colistin. This demonstrates the efficacy of TraDIS-*Xpress* for identifying antimicrobial targets and mechanisms of synergy with currently available antimicrobials. Furthering our understanding of cell responses to antimicrobial compounds may inform the design of future antimicrobial compounds and the synergistic relationships with novel antimicrobials, as a means of countering antimicrobial-resistant pathogens.

## Results & Discussion

### OPT-2U1 targets LPS biosynthesis

TraDIS-*Xpress* mutant libraries of *E. coli* and *S*. Typhimirium, containing over 800,000 and 500,000 unique transposon insertion mutants respectively, were used to investigate the genes required for survival in the presence of three analogues of OPT-2U1 (analogues 003, 009 and 010). Each analogue was tested at concentrations of 8, 16, 32 and 64 µg/mL, equivalent to ¼x, ½x, 1x and 2x the minimum inhibitory concentration (MIC) of the antimicrobial. Transposon insertion sites and frequencies were compared between cultures treated with the OPT-2U1 analogues and untreated controls to identify where transposon mutants or the modulation of gene expression (driven from the transposon-located outwards-transcribing promoter inserting upstream) have impaired or improved fitness under the test condition (supplementary figure 1). Analysis of the TraDIS-*Xpress* data cummulatively found 180 genes involved in survival across each of the three antimicrobial analogues in *E. coli* BW25113 and *S*. Typhimurium 14028*S* (supplementary table 1). These genes had roles in LPS biosynthesis, fatty acid biosynthesis, cell wall maintenance, cell division, efflux and respiration. From each of these pathways, 26 genes from *E. coli* and 5 genes from *S*. Typhimurium were then selected for validation of the TraDIS-*Xpress* data by investigating the susceptibility of gene deletion mutants to the OPT-2U1 analogues (supplementary table 1).

The most notable finding from the TraDIS-*Xpress* data was that increased expression of *lpxD* was beneficial for survival in the presence of each of the OPT-2U1 analogues. With increasing concentrations of each of the three analogues, more sequencing reads mapped upstream of *lpxD* in both *E. coli* (figure 1a) and *S*. Typhimurium (figure 1b) with the outward-transcribing promoter oriented to increase transcription of *lpxD*, indicating its expression was beneficial for survival in these conditions. There was upwards of a 9 log_2_-fold increase in the number of insertions upstream of *lpxD* in *E. coli* across the four concentrations of the three analogues of OPT-2U1, and a 2-14 log_2_-fold increase in *S*. Typhimurium, relative to the unstressed controls in each species (supplementary table 1). *LpxD* is an essential gene that encodes a UDP-3-O-(3-hydroxymyristoyl) glucosamine N-acyltransferase and is responsible for catalysing the third step of lipid IV_A_ biosynthesis, involved in making the lipopolysaccharide (LPS) core (Kelly et al., 1993). Because *lpxD* is an essential gene, its importance in the presence of an inhibitor would not normally be identified using conventional gene deletion or transposon mutagenesis techniques. This demonstrates the power of TraDIS-*Xpress* to assay the whole bacterial genome and identify potential targets of novel antimicrobial compounds accurately and reproducibly.

**Figure 1:**
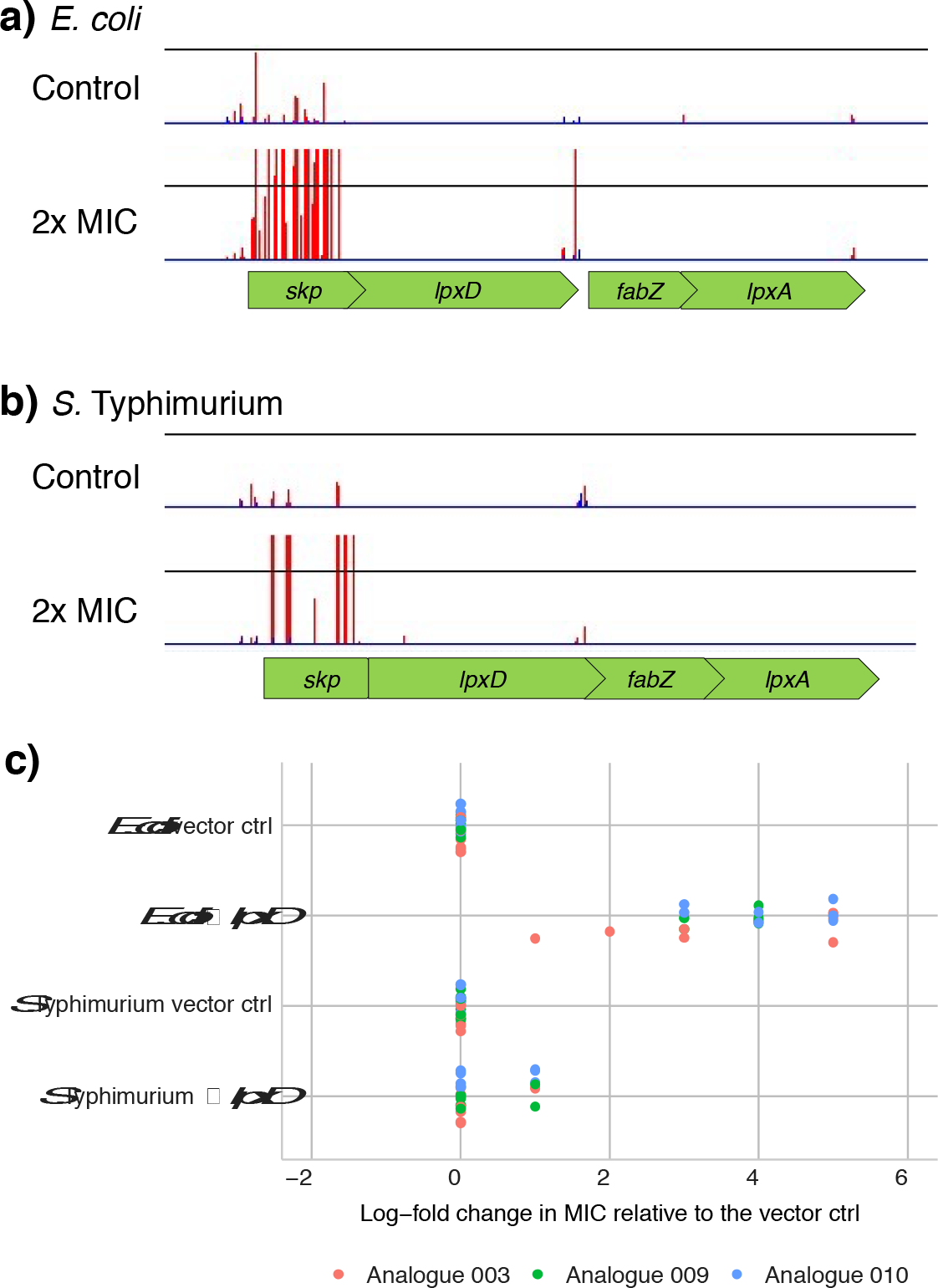
Increased expression of *lpxD* reduces susceptibility to OPT-2U1-003. **a)** Transposon insertion sites and frequencies around *lpxD* in *E. coli* and **b)** *S*. Typhimurium when treated with 2x the MIC of OPT-2U1 analogue 003 relative to unstressed controls. Reads were plotted with BioTraDIS in Artemis. Each vertical line indicates the location of a transposon insertion and the height indicates the approximate relative number of mutants at that site. Red lines mark where the transposon-located promoter is oriented left-to-right, and blue right-to-left. The Y-axes have been normalised for each locus to show relative differences in insert abundance between conditions. Experiments were performed in duplicate, but data from one only is presented due to the high consistency. **c)** Log_2_-fold change in MIC of the three analogues of OPT-2U1 in *E. coli* and *S*. Typhimurium overexpressing *lpxD* on a plasmid vector relative to an empty vector control in each species. The grey area indicates an experimental error of 1 log_2_-fold change from the wild type. Each point represents each of two technical and four biological replicates.

The effect of *lpxD* expression on antimicrobial susceptibility was confirmed by overexpressing the gene on a plasmid vector in both *E. coli* and *S*. Typhimurium. The native gene from each species was incorporated into a plasmid vector and introduced back into the parent strain. Overexpression of *lpxD* in this plasmid vector in *E. coli* resulted in a maximum MIC increase of 5 log_2_-fold for analogues 003 and 010, and 4 log_2_-fold for analogue 009, relative to a vector without *lpxD* (figure 1c). This confirmed the TraDIS-*Xpress* data, indicating that *lpxD* is a candidate antimicrobial target. However, overexpressing *lpxD* in *S*. Typhimurium did not confer a significant difference in susceptibility relative to the vector-only control or the parent strain without the vector (figure 1c). The *E. coli* K-12 strain used in this experiment is known to produce an incomplete LPS due to lack of O-antigen and *S*. Typhimurium 14028*S* produces a full LPS with a ‘smooth’ colony morphology. The difference in LPS composition between the two strains may result in the difference in susceptibility to OPT-2U1 analogues when *lpxD* is overexpressed.

### OPT-2U1 derivatives synergise with rifampicin and colistin, and this is affected by LPS composition

Each OPT-2U1 analogue acted synergistically with rifampicin or colistin to inhibit bacterial growth. At sub-MIC concentrations of the OPT-2U1 analogues, the MIC of rifampicin was reduced relative to OPT-free conditions in both *E. coli* and *S*. Typhimurium (figure 2a). However, synergy between OPT-2U1 and colistin was more pronounced in *S*. Typhimurium (figure 2b).

**Figure 2:**
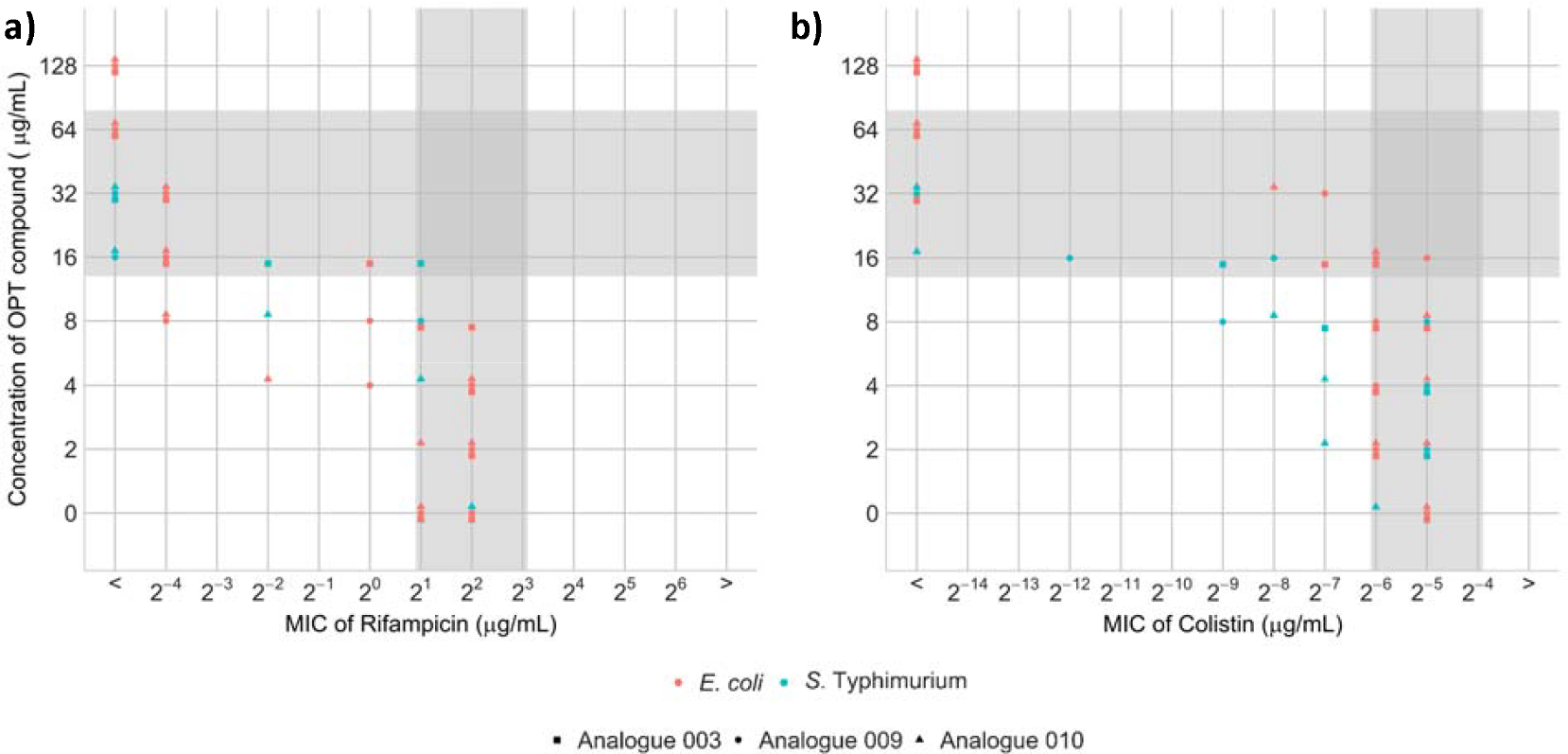
MICs of **a)** Rifampicin and **b)** Colistin in *E. coli* BW25113 and *S*. Typhimurium 14028*S* in the presence of sub-MIC concentrations of the three analogues of OPT-2U1 (003, 009 and 010). The grey area indicates an experimental error of 1 log_2_-fold change from the wild type. Points below and to the left of the grey experimental error ranges indicate the antibacterial combination acts synergistically. Each point represents each of two technical and two biological replicates.

We predicted the interspecies difference in synergy was due to LPS composition. Synergy between these antibiotics and OPT-2U1 was investigated in *S*. Typhimurium strains deficient in genes involved in LPS core (Δ*rfaB* and Δ*rfaK*) and O-antigen biosynthesis (Δ*rfbF*) and in *E. coli* with increased expression of lpxD (↑*lpxD*) involved in lipid IV_A_ biosynthesis (figure 3). Rifampicin is a large molecule that normally cannot cross the LPS unaided, so we investigated whether LPS disruption by OPT-2U1 caused synergy with rifampicin. Deletion of *rfaB*, but not *rfaK*, resulted in a reduced rifampicin MIC relative to wild type *S*. Typhimurium at sub-MIC concentrations of the analogues, despite both genes being involved in synthesis of the LPS core (figure 3a). Overexpression of *lpxD* on a plasmid vector in *E. coli* reduced the MIC of rifampicin relative to an empty vector control at sub-MIC concentrations of the analogues. When treated with sub-MIC concentrations of analogue 101, deletion of *rfbF* (involved in O-antigen biosynthesis of the LPS) resulted in 3 log_2_-fold reduction in the MIC of rifampicin in one replicate and a 2 log_2_-fold increase in two other replicates (figure 3a). This highlights the importance of investigating antibiotic synergies in all analogues across bacteria with different LPS compositions to identify the best candidates for progression.

**Figure 3:**
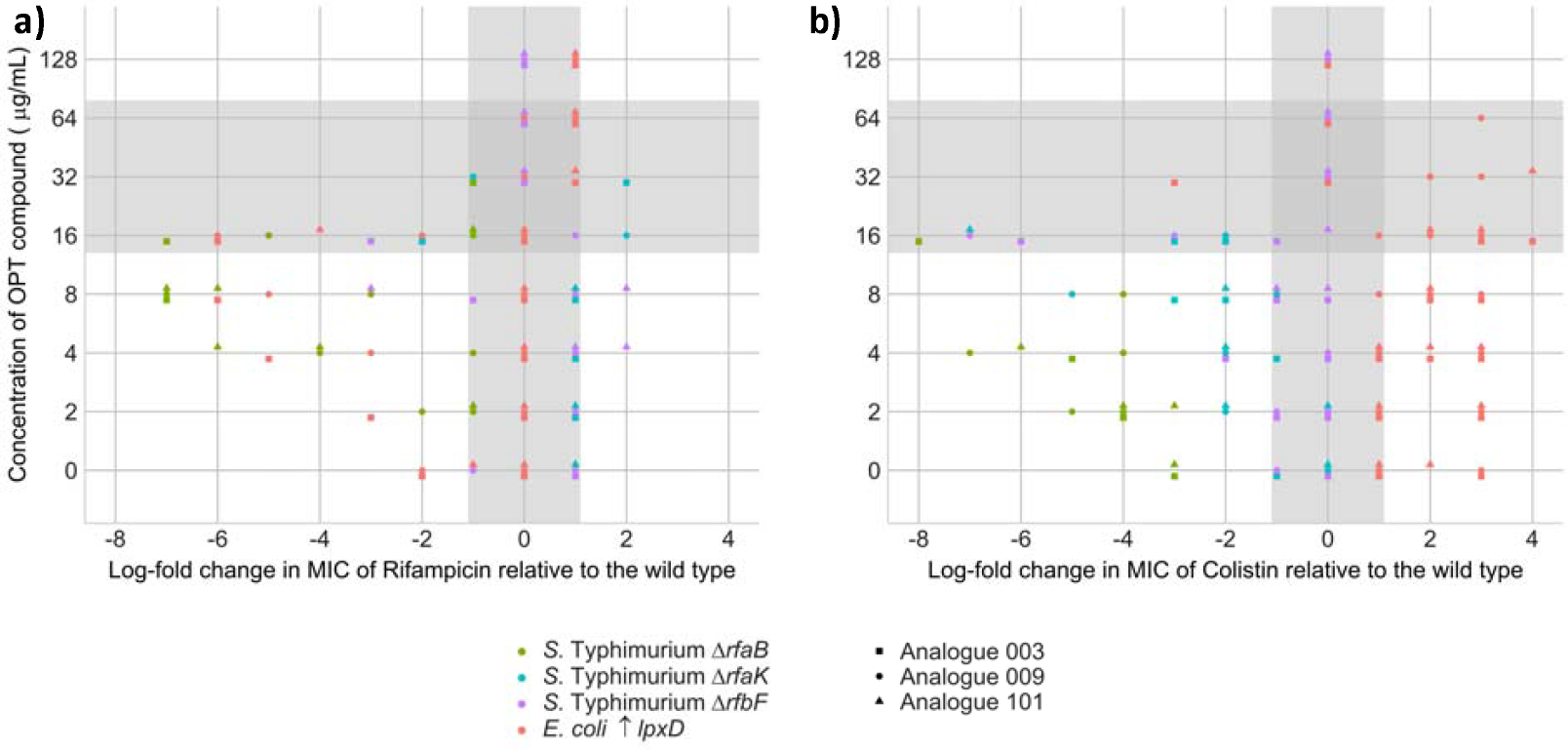
Log_2_-fold change in the MICs of **a)** Rifampicin and **b)** Colistin in the presence of sub-MIC concentrations of the three antimicrobial analogues of OPT-2U1 (003, 009 and 010) in *S*. Typhimurium gene deletion mutants Δ*rfaB*, Δ*rfaK*, Δ*rfbF* relative to the wild type and *E. coli* overexpressing *lpxD* (↑*lpxD*) relative to an empty vector control. The grey area indicates an experimental error of 1 log_2_-fold change from the wild type. Points below and to the left of the grey area indicate reduced growth of the mutants relative to the wild type in the presence of the drug combination. Points below and to the right of the grey area indicate increased growth in these conditions. Each point represents each of two technical and two biological replicates.

Colistin acts directly on the cell envelope, so we hypothesised that LPS composition would affect the synergy between colistin and the OPT-2U1 derivatives. Deletion mutants of genes involved in LPS core (*rfaB* and *rfaK*) and O-antigen (*rfbF*) biosynthesis had a lower MIC of colistin relative to wild type S. Typhimurium at sub-MIC concentrations of the analogues (figure 3b). Increased expression of *lpxD* on a plasmid vector in *E. coli* resulted in a 2 to 3 log_2_-fold increase in colistin MIC relative to a control harbouring an empty vector at sub-MIC concentrations of OPT-2U1 (figure 3b). This suggests disruption to LPS synthesis leads to increased colistin susceptibility, and increased expression of *lpxD* may increase lipid IV_A_ synthesis, reducing colistin susceptibility. This supports our hypothesis that differences in LPS composition between *E. coli* and *S*. Typhimurium may cause the difference seen in the synergistic relationship between OPT-2U1 analogues and colistin for each species.

### ATP synthesis and the BasSR signalling system affect synergy between the OPT-2U1 and colistin

Analysis of the TraDIS-*Xpress* data revealed more insertions across the operon encoding ATP synthase (*atpIBEFHAGD*) in *S*. Typhimurium in conditions treated with the OPT-2U1 analogue 010 relative to the untreated control (figure 4a). An *atpB* deletion mutant in *E. coli* did not significantly affect susceptibility to any of the OPT-2U1 analogues (figure 4b), but did affect the synergy between OPT-2U1 and colistin: there was a 2-to-3 log_2_-fold reduction in the MIC of colistin at sub-MIC concentrations of the OPT-2U1 analogues in the Δ*atpB* deletion mutant relative to the wild type (figure 4c). Deletion of genes encoding ATP synthase have previously been found to increase susceptibility to colistin (Yasir et al., 2022), and we found the addition of OPT-2U1 further increased colistin susceptibility in *E. coli*.

**Figure 4:**
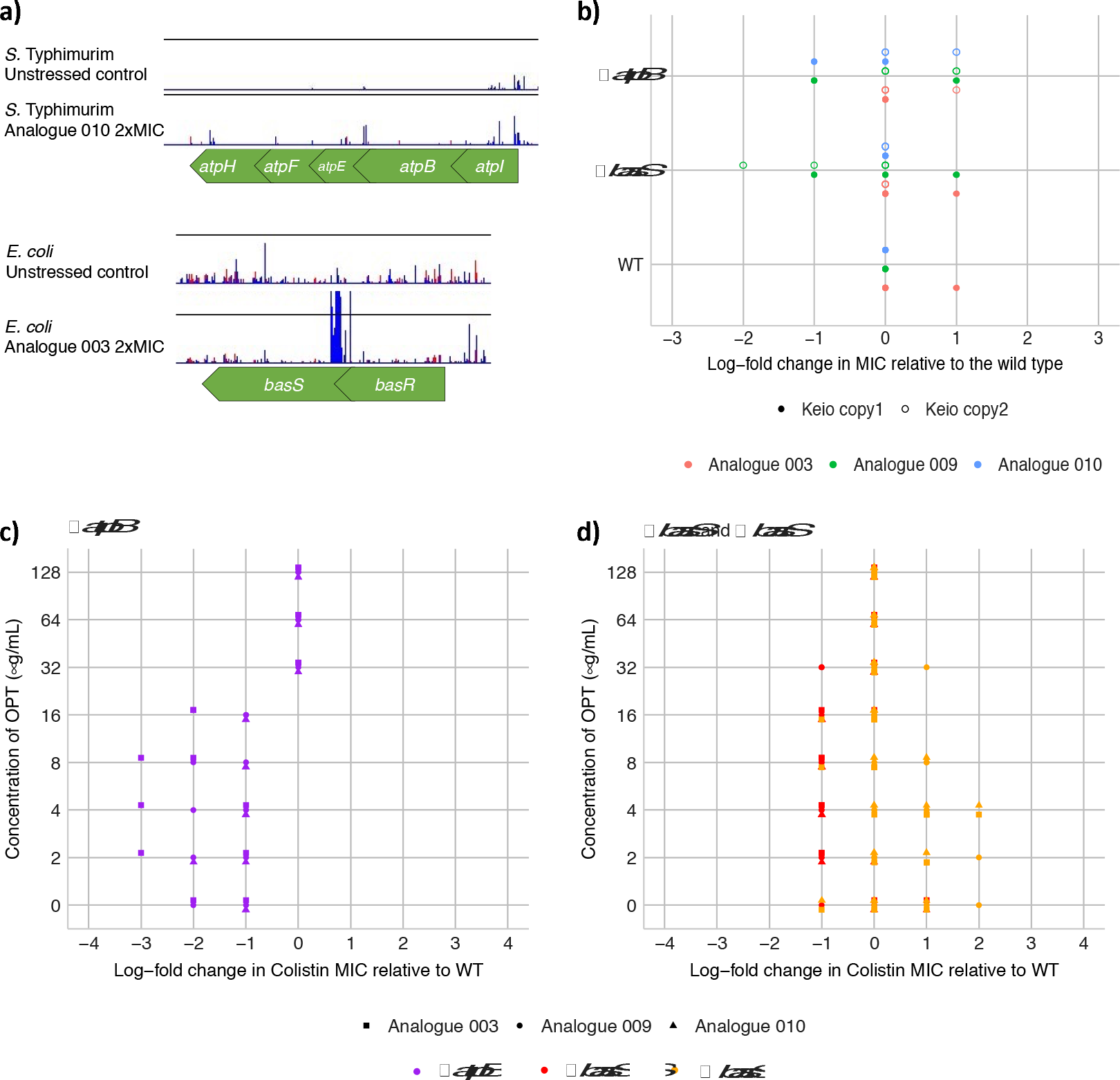
**a)** Transposon insertion sites and frequencies around the *atp* operon in *S*. Typhimurium and *basS* in *E. coli* when treated with the OPT-2U1 analogues 010 and 003 respectively, relative to unstressed controls. Reads were plotted with BioTraDIS in Artemis. Each vertical line indicates the location of a transposon insertion and the height indicates the approximate relative number of mutants at that site. Red lines mark where the transposon-located promoter is oriented left-to-right, and blue right-to-left. The Y-axes have been normalised for each locus to show relative differences in insert abundance between conditions. Experiments were performed in duplicate, but data from one only is presented due to the high consistency. **b)** Log_2_-fold change in the MIC of the three OPT-2U1 analogues (003, 009 and 010) in gene deletion mutants relative to wild type *E. coli*. The Keio collection contains two copies of each gene deletion construct, which have been presented separately. **c)** Log_2_-fold change in the MIC of colistin at sub-MIC concentrations of the three analogues relative to wild type *E. coli*, following deletion of *atpB* (Δ*atpB*), **c)** *basS* (Δ*basS*) and overexpression of *basS* (↑*basS*) relative to an empty vector control. For all plots, the grey area indicates an experimental error of 1 log_2_-fold change from the wild type. In panels c and d, points below and to the left of the grey area indicate reduced growth of the mutants relative to the wild type in the presence of the drug combination. Points below and to the right of the grey area indicate increased growth in these conditions. Each point represents each of two technical and two biological replicates.

We found overexpression of *basS* to be beneficial for the survival of *E. coli* in the presence of OPT-2U1 analogues 003 and 009. Analysis of the TraDIS-*Xpress* data shows more insertions upstream of *basS* in conditions treated with these analogues relative to the untreated control (figure 4a). The BasSR signalling system regulates genes involved in membrane structure, membrane function and stress responses and has also been implicated in *E. coli* susceptibility to colistin (Knopp et al., 2021, Yasir et al., 2022). We investigated whether modulating the expression of *basS* affected synergy between OPT-2U1 and colistin in *E. coli*: a deletion mutant of *basS* showed in a 2 log_2_-fold reduction in MIC of OPT-2U1 analogue 009 (figure 4b) relative to the wild type, and did not affect the synergy between any OPT-2U1 analogue and colistin (figure 4d). Increased expression of *basS* on a plasmid vector resulted in a 2 log_2_-fold increase in the MIC of colistin at sub-MIC concentrations of all OPT-2U1 analogues in *E. coli* relative to the vector control (figure 4d). This warrants further research into the regulatory pathways through which *basS* affects the synergy between OPT-2U1 and colistin.

## Conclusions

Determining the mechanism of action of a novel antimicrobial is a crucial step for its progression through the antimicrobial development pipeline. We used TraDIS-*Xpress*, a massively parallel high throughput genome-wide screen, to determine the mechanism of action of three analogues of a novel antimicrobial compound, OPT-2U1 in both *E. coli* and *S*. Typhimurium. We propose the main target of this antimicrobial is likely to be *lpxD*, involved in synthesis of the lipid IV_A_ core of the LPS component of the cell’s outer membrane. LPS biosynthesis is an attractive target for antibiotic development and *lpxD* has previously been shown to be a potentially valuable target for inhibitor development with chemically tractable binding sites which are conserved across Enterobacteriaceae (Ma et al., 2020, Sabnis and Edwards, 2023). As well as the proposed primary target, other genes involved in LPS biosynthesis, membrane maintenance and efflux activity affected susceptibility to OPT-2U1 in both species tested (Figure 5). Synergies between the OPT-2U1 analogues combined with rifampicin or colistin were identified, but these varied between species and may be a result of differences in LPS composition between the strains tested. Genes encoding ATP synthase and the BasSR two component signalling system affected the synergy between OPT-2U1 and colistin, most likely through their effects on membrane structure and function. This further substantiates our findings of the importance of LPS and outer envelope integrity in the activity of OPT-2U1 (figure 5). This work shows OPT-2U1 analogues have promise for continued development as they target a conserved and essential structural component of Enterobacteriaceae and we identify possible synergies which could be exploited in formulations to increase efficacy and reduce the likelihood of resistance emergence. TraDIS-*Xpress* proved a rapid and sensitive tool that identified the likely mechanism of action of a novel antimicrobial, as well as the wider landscape of other genes which interact with these analogues. The ability of this method to assay essential genes makes it valuable as a tool for drug discovery.

**Figure 5:**
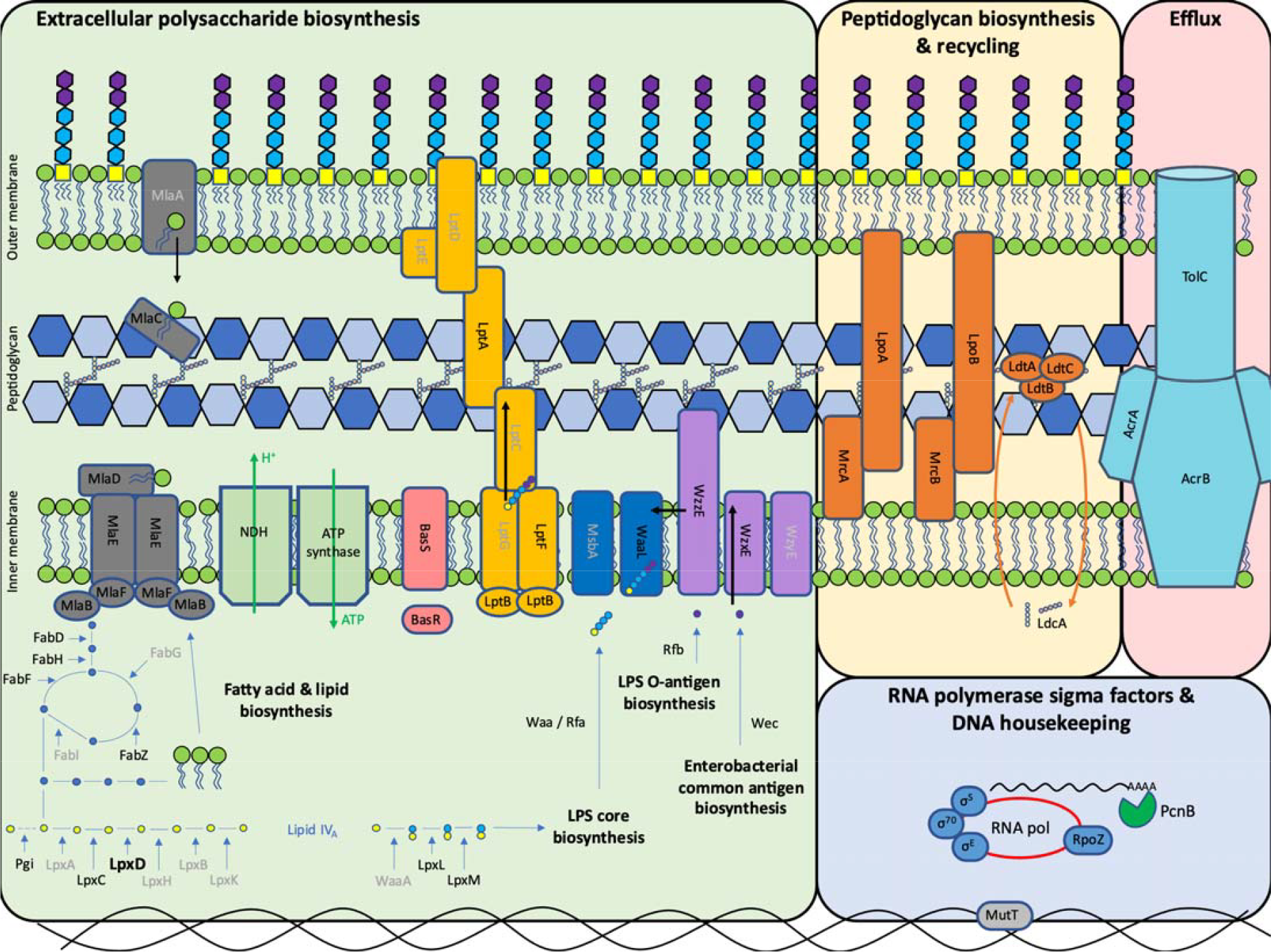
Cellular overview of the pathways and proteins encoded by genes that affect susceptibility to the novel OPT-2U1 analogues, shown here in black text. Genes encoded by proteins in grey text were not identified by TraDIS-*Xpress* to affect susceptibility but are included in the diagram for context.

## Methods

### Transposon mutant libraries and growth conditions

The *E. coli* BW25113 transposon mutant library used in this study was described by Yasir et al. (2020), and the *S*. Typhimurium 14028*S* library was described by Holden et al. (2022). The *S*. Typhimurium mutant library contains a chromosomal copy of *lacI* to facilitate control of the transposon-located *tac* promoter, which was achieved with 1 mM IPTG in both *E. coli* and *S*. Typhimurium libraries. For each condition, approximately 10^7^ CFU/mL of the transposon mutant library was added to 10 mL LB broth supplemented with the corresponding concentration of antimicrobial. Each of the three analogues of OPT-2U1 were investigated at 8, 16, 32 and 64 µg/mL, equivalent to ¼x, ½x, 1x and 2x the MIC. These were incubated for 24 hours at 37 □C with shaking at 200 rpm. Following this, 1 mL of each culture was centrifuged to form a bacterial pellet. DNA was extracted from these pellets following the protocol outlined by Trampari et al. (2021).

### Sequencing Library Preparation and Informatics

Genomic DNA concentrations were adjusted to 11.1 ng/µL and tagmented using a MuSeek DNA fragment library preparation kit (ThermoFisher). Fragmented DNA was purified with AMPure XP beads (Beckman Coulter). DNA was amplified using primers customised for the tagmented ends of the DNA and biotinylated primers specific to transposon. Following another purification step, biotinylated fragmented DNA was incubated for 4 hours with streptavidin beads (Dynabeads® kilobaseBINDER™, Invitrogen) to capture only DNA fragments containing the transposon. A second PCR step using DNA bound to the beads used barcoded sequencing primers specific to the tagmented ends and to the transposon. Beads were removed from the reaction with a magnet and DNA was purified and size-selected using AMPure beads. Fragment length was quantified using a Tapestation (Aligent) and sequenced on a NextSeq500 using the NextSeq 500/550 High Output Kit v2.5 with 75 cycles. Fastq files were aligned to the *E. coli* BW25113 reference genome (CP009273) and *S*. Typhimurium 14028*S* reference genome (CP001363, modified to include chromosomally integrated *lacIZ*) using BioTraDIS (version 1.4.3) (Barquist et al., 2016). Significant differences (*p* < 0.05, after correction for false discovery) in insertion frequencies between the unstressed controls and treated conditions were found using the tradis_comparison.R command (part of the BioTraDIS toolkit) and AlbaTraDIS (version 1.0.1) (Page et al., 2019).

### Construction of gene deletion mutants and expression vectors

Gene deletion mutants in *E. coli* were taken from the Keio collection (Baba et al., 2006). There are two copies of each gene deletion construct in the Keio collection, and these have been analysed separately to account for cloning errors between the copies. Gene deletion mutants in *S*. Typhimurium were made following the Gene Doctoring protocol (Lee et al., 2009) using vectors assembled using Golden gate assembly (Thomson et al., 2020). Overexpression vectors were also made using Golden gate assembly. *LpxD* was inserted into the IPTG-inducible vector pJMA*t5*, however leaky expression from the *lac* promoter prevented the assembly of *basS* into this vector, so *basS* was inserted into the rhamnose-inducible vector pJMA*rha* (both vectors kindly gifted by Nicholas Thomson). For each construct, the gene of interest was amplified from the parent organism using primers with BsaI cut sites incorporated at the 5’-ends. Amplified DNA fragments encoding the relevant gene were combined with the relevant vector plus BSA, T4 ligase and buffer and the BsaI restriction enzyme, as described by Thomson et al. (2020). This mixture underwent 30 cycles of 37 □C for 3 minutes and 16 □C for 4 minutes, followed by 50 □C for 5 minutes and 80 □C for 5 minutes. Chemically competent cells were transformed with 2 µL of these assemblies and spread on LB agar containing 50 µg/mL kanamycin. Following growth at 37°C, resulting colonies were screened by PCR and validated by Sanger sequencing to identify the correct recombinant plasmid.

### Antimicrobial susceptibility and synergies

Antimicrobial susceptibility was measured using a broth dilution method following EUCAST guidelines (EUCAST, 2021). Synergies between antibiotics were investigated using a checkerboard assay modified from the broth dilution method, where the concentration of one antimicrobial decreases across the plate left-to-right and the second antimicrobial decreases in concentration down the plate. The MIC of rifampicin or colistin was determined at each sub-MIC concentration of the OPT-2U1 analogue. This was repeated with two biological and two technical replicates. For assays of strains carrying expression vectors, increased expression of *lpxD* was induced with 1 mM IPTG and increased expression of *basS* was induced with 0.02% rhamnose.

## Competing interests

AM, JG and TR are founding shareholders in Oppilotech Ltd. All other authors declare no competing interests.

## Funding Information

ERH, ES, AM, JG and MG were supported by Innovate UK project number 104991. The authors gratefully acknowledge the support of the Biotechnology and Biological Sciences Research Council (BBSRC): ERH, MY, AKT, MAW and MG were supported by the BBSRC Institute Strategic Programme Microbes in the Food Chain BB/R012504/1 and its constituent project BBS/E/F/000PR10349.

## Notes

https://www.ebi.ac.uk/biostudies/arrayexpress/studies/E-MTAB-13250

