## Supplementary figure 1 for "Mechanisms of action and synergies of a novel lipid IV_A_ biosynthesis inhibitor"

**Supplementary figure 1:** Insertion frequency per gene for the *E. coli* transposon mutant library cultured under four concentrations (¼x MIC, ½x MIC, 1x MIC and 2x MIC) of each of the three analogues of OPT-2U1: **a)** OPT003, **b)** OPT009 and **c)** OPT010. The *S*. Typhimurium transposon mutant libraries were also exposed to **d)** OPT003, **e)** OPT009 and **f)** OPT010. Black points represent the insertion frequency per gene for each replicate to show variation between the replicates, and coloured points show the insertion frequency per gene of the control against the antimicrobial in each condition.

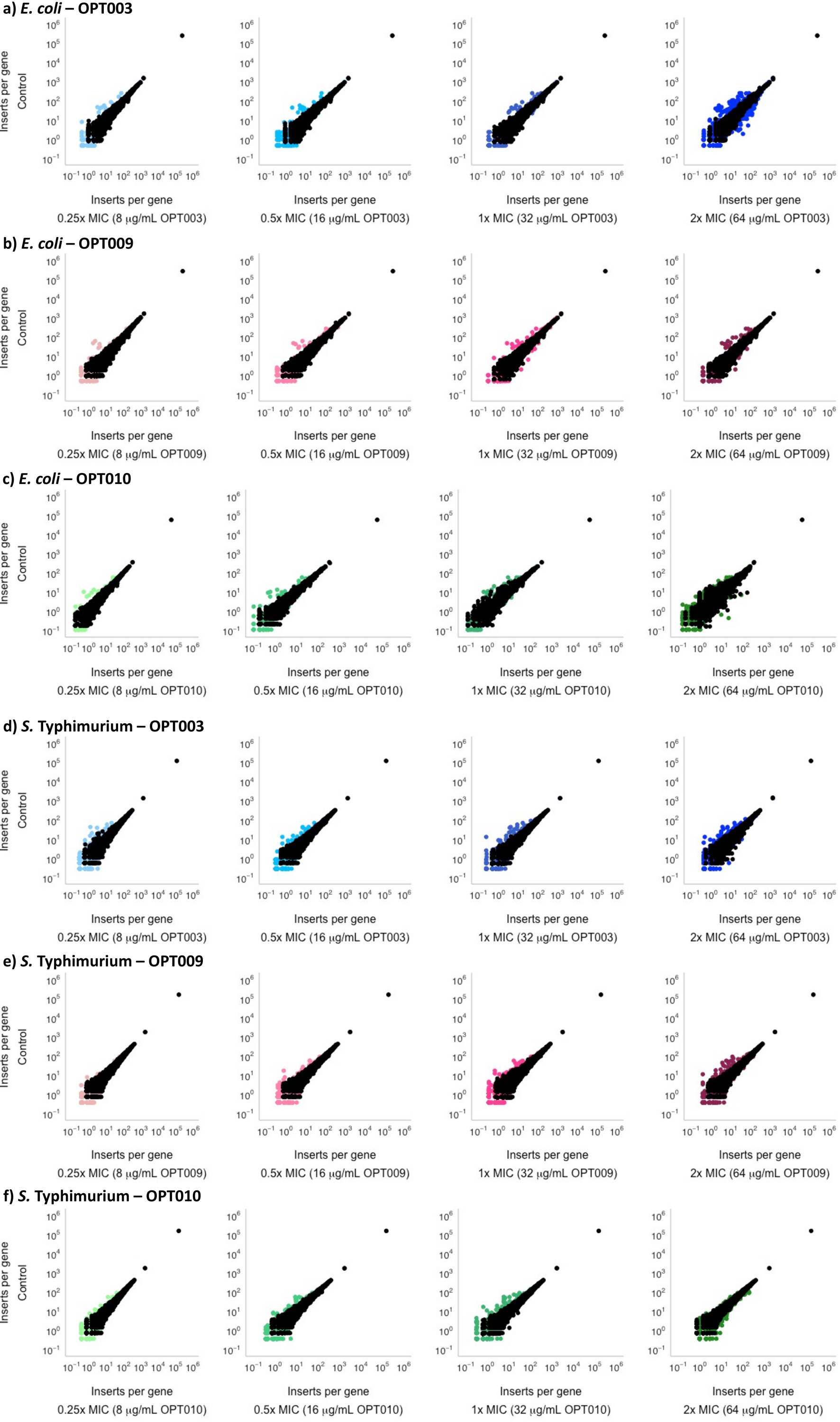
